# Evidence of SARS-CoV2 entry protein ACE2 in the human nose and olfactory bulb

**DOI:** 10.1101/2020.07.15.204602

**Authors:** M. Klingenstein, S. Klingenstein, P.H. Neckel, A. F. Mack, A. Wagner, A. Kleger, S. Liebau, A. Milazzo

**Author notes:** equal contribution (M. Klingenstein, S. Klingenstein).

## Abstract

Usually, pandemic COVID-19 disease, caused by SARS-CoV2, presents with mild respiratory symptoms such as fever, cough but frequently also with anosmia and neurological symptom. Virus-cell fusion is mediated by Angiotensin-Converting Enzyme 2 (ACE2) and Transmembrane Serine Protease 2 (TMPRSS2) with their organ expression pattern determining viral tropism. Clinical presentation suggests rapid viral dissemination to central nervous system leading frequently to severe symptoms including viral meningitis. Here, we provide a comprehensive expression landscape of ACE2 and TMPRSS2 proteins across human, post-mortem nasal and olfactory tissue. Sagittal sections through the human nose complemented with immunolabelling of respective cell types represent different anatomically defined regions including olfactory epithelium, respiratory epithelium of the nasal conchae and the paranasal sinuses along with the hardly accessible human olfactory bulb. ACE2 can be detected in the olfactory epithelium, as well as in the respiratory epithelium of the nasal septum, the nasal conchae and the paranasal sinuses. ACE2 is located in the sustentacular cells and in the glandular cells in the olfactory epithelium, as well as in the basal cells, glandular cells and epithelial cells of the respiratory epithelium. Intriguingly, ACE2 is not expressed in mature or immature olfactory receptor neurons and basal cells in the olfactory epithelium. Similarly ACE2 is not localized in the olfactory receptor neurons albeit the olfactory bulb is positive. Vice versa, TMPRSS2 can also be detected in the sustentacular cells and the glandular cells of the olfactory epithelium.

Our findings provide the basic anatomical evidence for the expression of ACE2 and TMPRSS2 in the human nose, olfactory epithelium and olfactory bulb. Thus, they are substantial for future studies that aim to elucidate the symptom of SARS-CoV2 induced anosmia of via the olfactory pathway.

## INTRODUCTION

The Coronavirus Disease 2019 (COVID-19) emerged from East Asia and quickly spread all over the world reaching a pandemic scale [1]. The infiltration of the virus SARS-CoV2 into different cell types leads to different symptoms with varying severity. According to clinical studies worldwide, the most prevalent symptoms are fever, cough, fatigue, headache, dyspnea, sputum production, arthralgia, diarrhea, rhinorrhea, and sore throat [2, 3]. Since many patients also report olfactory and gustatory dysfunctions, these symptoms are considered typical of the SARS-CoV2 infection [4, 5]. There are many steps involved in the perception of smell where the infection with SARS-CoV2 could potentially be the cause of anosmia, starting with the transport of the odorants to the receptors in the olfactory neurons extending to the signal transduction to different olfactory cortex areas.

In the olfactory epithelium, a variety of histological target structures, including the olfactory receptor neurons (ORN) with their ensheathed axons, the sustentacular, microvillar or glandular cells could serve as a viral target and therefore influencing olfactory function. Another area of viral attack could be the olfactory bulb. Here, the fila olfactoria or projection neurons in different layers of the olfactory bulb could be targeted by the virus, causing disruption in olfactory perception.

Infection of host cells with SARS-CoV-2 is preceded by a complex process of virus attaching, receptor recognition and proteolytic cleavage of the transmembrane spike glycoprotein to promote virus-cell fusion mediated by Angiotensin-Converting Enzyme 2 (ACE2) and Transmembrane Serine Protease 2 (TMPRSS2) [6].

Up to date, there is no clear evidence which cell types of the olfactory and respiratory epithelium express ACE2 and TMPRSS2. Transcriptional but also data from murine and human olfactory tissue show as of now report conflicting data [7-10]. In addition, the reports on localization of the viral entry proteins in different cell types from the epithelia vary [7, 8]. Protein expression was found in sustentacular cells in all publications, but only one paper claims a low ACE2 expression in ORN [8]. TMPRSS2 was found in murine and human respiratory epithelium and murine olfactory epithelium [7]. Similarly, experiments performed in the murine olfactory bulb showed partly contradictory results for ACE2 staining. Until now, there are no protein verifications in human olfactory bulb due to its difficult accessibility.

Here, we employ a unique human post-mortem tissue resource to thoroughly study the ACE2 and TMPRSS2 protein expression patterns in the human olfactory system, including olfactory epithelium, respiratory epithelium of the nasal septum, the nasal conchae and the paranasal sinuses as well as the olfactory bulb. These findings will help to explain the symptom anosmia as well as frequent dissemination to the central nervous system in COVID-19 patients and give a starting point to further investigations how SARS-CoV2 can affect the olfactory system.

## RESULTS

### Anatomical regions analyzed in human post-mortem nasal and olfactory tissue

High ACE2 and TMPRSS2 expressions have been documented in the respiratory, gastrointestinal, and reproductive system [11, 12]. Focusing on the upper respiratory tract, increased ACE2 and TMPRSS2 are found in the respiratory and olfactory mucosa. First, a frontal section of human post-mortem tissue through the nose with distinct anatomical regions (Figure **1A**) is examined. The olfactory epithelium (IV) is located at the roof of the nasal cavity between the nasal septum and the superior nasal conchae. The paranasal sinuses, in particular the cellulae ethmoidales (III), are located next to the nasal turbinate. In addition, the nasal septum (I), the superior nasal conchae, and the intermediate nasal conchae (II), all of which covered with respiratory epithelium, are present in this section (Figure **1B**). With Hematoxylin-Eosin staining, these different regions of the nasal cavity were identified and visualized (Figure **1C**; **2B**; **3B**). Overview of ACE2 and TUBB3 immunolabelling in post-mortem nasal tissue (Figure **1D**). TUBB3, a marker for mature and immature olfactory sensory neurons allows distinguishing the olfactory epithelium from the other non-neural areas within the nose specimen (Figure **1D**).

**Figure 1:**
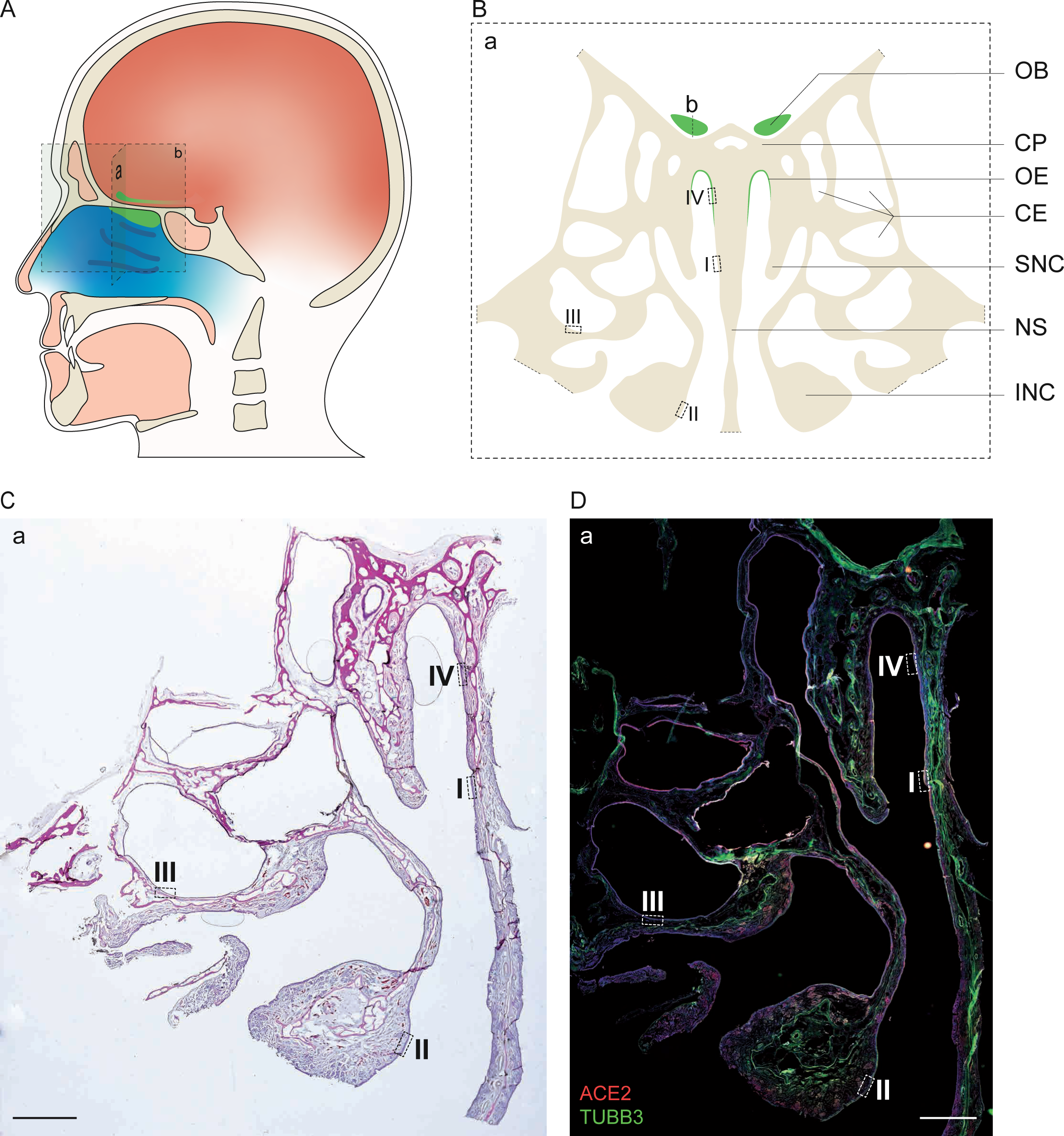
(A) Schematic illustration of a human head. Frontal section highlights the nose specimen (a). Sagittal view of the olfactory bulb section (b). Olfactory epithelium and olfactory bulb, both marked in green, are located at the upper nasal cavity and above the cribriform plate. (B) Schematic representation of the nose specimen from the frontal section through the head, shown in (a). The olfactory epithelium (OE) and the olfactory bulb (OB) are shown in green. (b) Section plane for better comprehensibility of the olfactory bulb. Dashed boxes show the section of the respiratory epithelium of the nasal septum (NS) (I), the intermediate nasal conchae (INC) (II), the cellulae ethmoidales (CE) (III) and the olfactory epithelium (OE) (IV). Other abbreviations: CP: Cribriform plate, SNC: Superior nasal conchae. (C). HE staining of the right half of nose specimen. Dashed boxes show the same areas found in the schematic picture in (B). (D) Immunofluorescence staining of the right half of human nose specimen. TUBB3 is shown in green, ACE2 in red, nuclei in blue. Dashed boxes show the same areas found in the schematic picture in (B). More details for area I, II, III of the respiratory epithelium can be found in Figure 2.More details for area IV, olfactory epithelium can be found in Figure 3. Scale bar: 2,5 mm.

**Figure 2:**
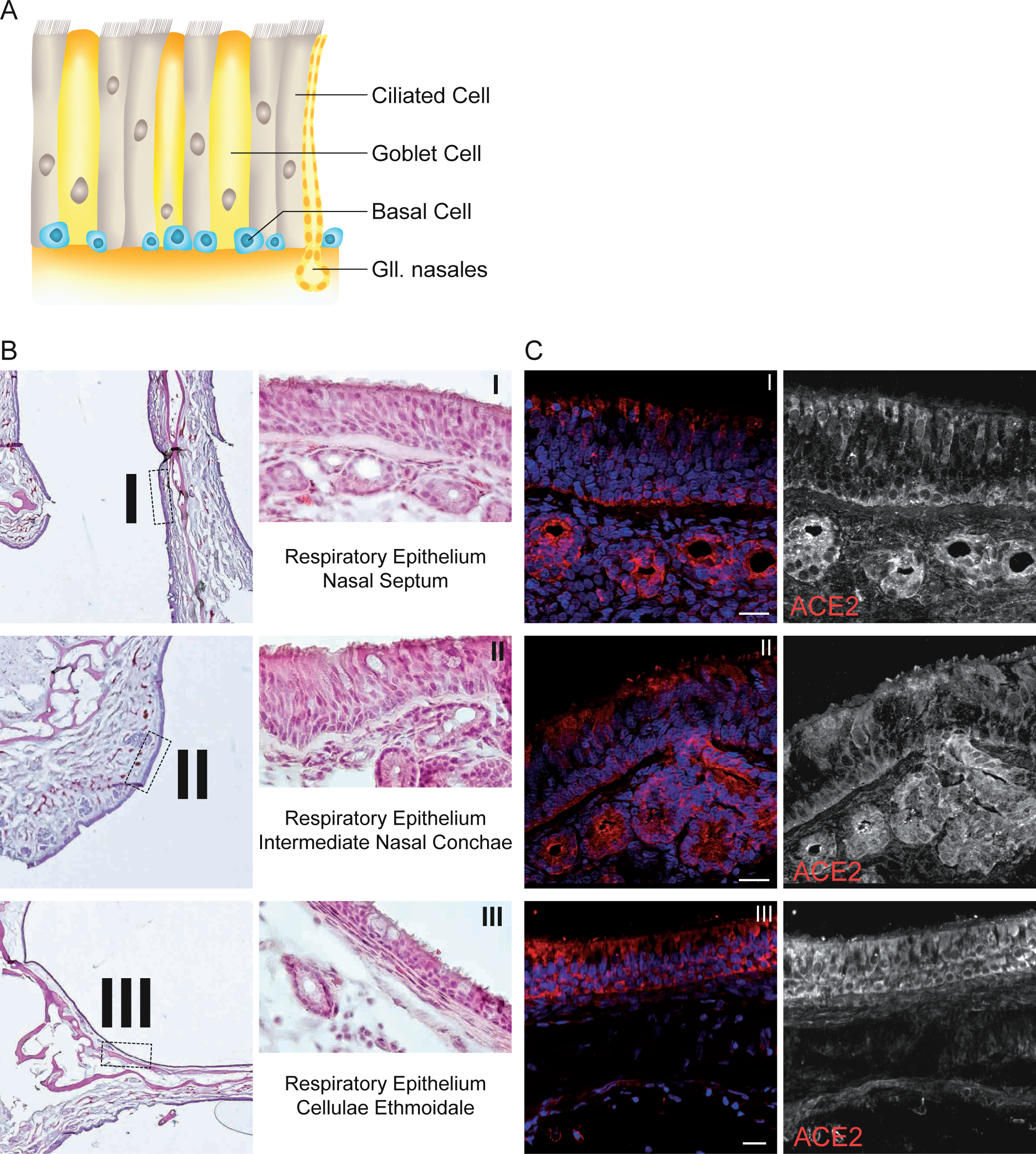
(A) Schematic illustration of the pseudostratified respiratory epithelium with ciliated cells, goblet cells, basal cells and submucosal Gll. nasales. (B) HE stainings of the respiratory epithelium in the area of the nasal septum (I), the intermediate nasal conchae (II), and the cellulae ethmoidales (III). (C) Immunofluorescent stainings of the respiratory epithelium verified by absent TUBB3 expression in the epithelium from dashed boxes I, II and III. TUBB3 is a highly specific marker for mature and immature ORN. ACE2 (red) positive basal cells of the respiratory epithelium of the nasal septum as well as positive apical staining of respiratory epithelial cells and Gll. nasals (I). ACE2 (red) protein expression is located in the ciliated epithelium cells and the basal cells of the respiratory epithelium of intermediate nasal conchae as well as in the underlying Gll. nasals (II). ACE2 (red) expression in the cellulae ethmoidales can be found in epithelial cells and the basal cells (III). Nuclei are shown in blue, scale bar: 20 µm.

**Figure 3:**
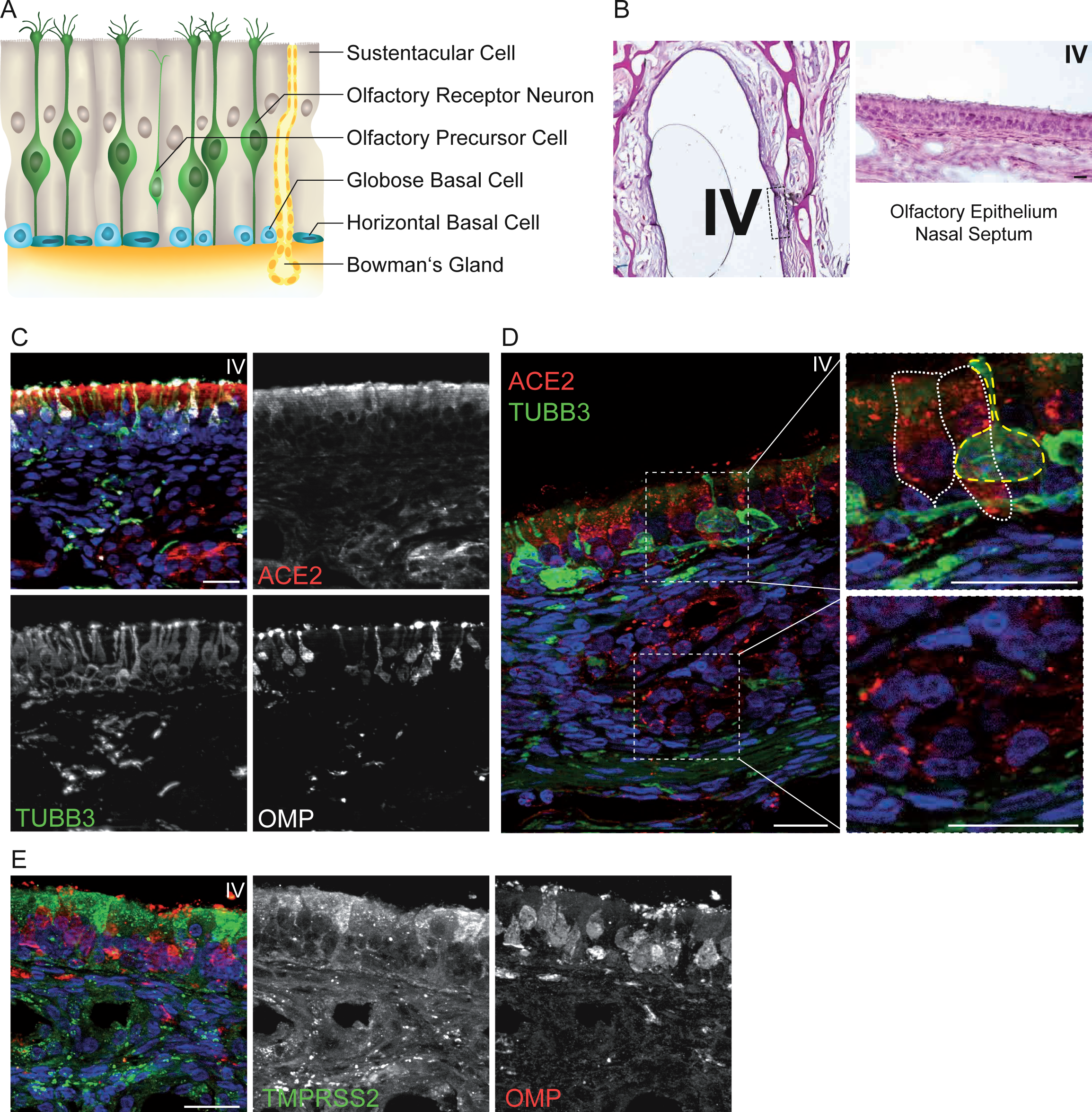
(A) Schematic structure of human olfactory epithelium. Mature olfactory receptor neurons are highlighted in green and are surrounded by sustentacular cells. The sensory neurons arise via an olfactory precursor step from basal cells. There are two types of basal cells: Horizontal and globose basal cells. Bowman’s glands are secretory glands which are found exclusively in the olfactory epithelium. (B) HE staining of the olfactory epithelium in the area of the nasal septum (IV). (C) Immunofluorescent staining of the olfactory epithelium from dashed box (IV). Merged picture shows ACE2 in red, OMP, a marker for mature ORN, in white and TUBB3, a marker for mature and immature ORN, in green. Positive ACE2 staining is exclusively located to the region of the sustentacular cells, no appearance in basal cells or co-localization with OMP or TUBB3. (D) Enlarged picture from the olfactory epithelium and mucosa showing ACE2 positive sustentacular cells and positive Bowman’s glands. Upper dashed box shows enlarged picture of ACE2 positive sustentacular cell, marked with dashed white lines, but negative for TUBB3. TUBB3 positive olfactory sensory neuron marked with dashed yellow lines. Lower dashed box shows ACE2 positive staining in Bowman’s glands in the olfactory submucosa. (E) Immunofluorescent picture with TMPRSS2 (green) and OMP (red) expression in the olfactory epithelium (IV). TMPRSS2 is mainly located in the sustentacular cells and shows a minor expression in Bowman’s glands, but no expression in basal cell or OMP-positive mature ORN. Nuclei are shown in blue, scale bar: 20 µm.

### Identification of ACE2 in different areas of the respiratory epithelium

Schematic illustration and Hematoxylin-Eosin staining shows the anatomical stratification of the analyzed regions of the nasal cavity (Figure **1C**; **2A; 2B**). The different areas of the respiratory epithelium, namely from the nasal septum (I), the intermediate nasal conchae (II) and the cellulae ethmoidales (III) show pseudostratified ciliated epithelial cells arising from a basal cell layer. Mucus-producing goblet cells can be found in between in different size and number (Figure **2B**). In the respiratory epithelium of the nasal septum, the intermediate nasal conchae and the paranasal sinus, ACE2 expression was located in epithelial cells as well as in basal cells. High expression was found in the Gll. nasales underneath the respiratory epithelium (Figure **2C**).

### Identification of ACE2 and TMPRSS2 in the olfactory epithelium

Schematic illustration of the olfactory epithelium (Figure **3A**) and histological staining of the same area (Figure **3B**). The Olfactory epithelium includes several distinct cell types, the most important of these are primary sensory neurons which are embedded between the supporting cells. These sustentacular cells provide mechanical strength to the epithelium, generate the olfactory binding protein, and support the other cells with nutrients [13]. Additionally, sustentacular cells are responsible for the maintenance of the ion and water balance within the olfactory epithelium [14]. ORN arise from basal stem cells via immature and intermediate steps [15]. Moreover, microvillar cells and excretory ducts from the specialized Bowman’s glands beneath the epithelium lie scattered in the epithelium [16, 17]. In addition, the data confirmed the expression of ACE2 in the olfactory epithelium (Figure **3C,D**). The ACE2 staining was mainly located in the supporting cells and could not be found in basal cells, immature ORN, marked with TUBB3 nor in mature ORN, marked with olfactory marker protein (OMP) (Figure **3C**). In the underlying lamina propria high concentrations of ACE2 were found in the Bowman’s gland cells. ACE2 expression is mainly limited to the sustentacular cells and shows no co-localization with adjacent TUBB3 positive sensory neurons (Figure **3D**). TMPRSS2 was also expressed in the supporting cells of the olfactory epithelium and in the glandular cells of both epithelia, but no expression in the other cell types (Figure **3E**).

### Identification of ACE2 in the olfactory bulb

To complete our view of ACE2 expression in the olfactory system, stainings of the human olfactory bulb were performed. Axons from the ORN pervade through the lamina cribrosa to the olfactory bulb. Here, the first synaptic interconnections to the downstream secondary neurons appear [18]. These defined interconnections are located in six distinct layers of the olfactory bulb (Figure **4A**,**C**). With Hematoxylin-Eosin staining, these different regions of the olfactory bulb were identified and visualized (Figure **4B**). The processed information projects along the olfactory tract to different regions of the olfactory cortex [19]. The hallmark feature of the outer nerve layer is the ensheated fila olfactoria coming from the olfactory epithelium, penetrate the bone plate and enter the olfactory bulb. Different cell types and their interconnections, like interneurons and projection neurons are located in deeper layers of the olfactory bulb and lead to processed and fine-tuned sensory information (Figure **4A**) [20]. The olfactory bulb has been analyzed on RNA level and on protein level with mouse tissue so far [8, 9]. In the human olfactory bulb, ACE2 could be found widely distributed, with high expression in the glomerular layer (Figure **4D**). Faint stainings of ACE2 in the mitral cell layer was also detected. However, no co-localization of ACE2 and OMP or TUBB3 could be detected, suggesting that ACE2 is not expressed in the axons of the ORN or other neurons (Figure **4D**).

**Figure 4:**
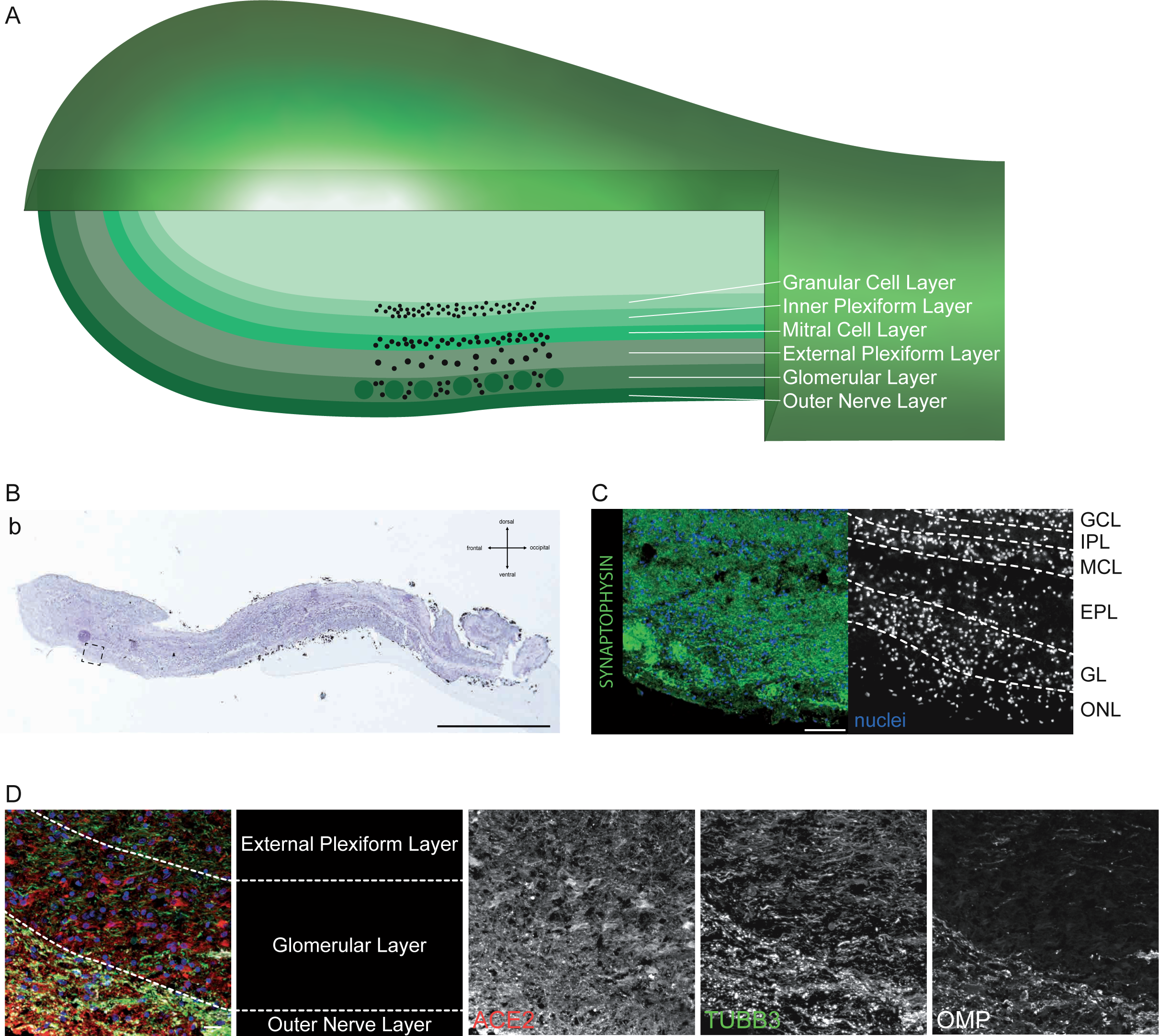
(A) Schematic illustration of the human olfactory bulb with its layers: Undermost is the outer nerve layer with the axons of the olfactory receptor neurons followed by the glomerular layer with olfactory glomerula connecting the axons with interneurons. Synaptic processing between the glomerular layer and the mitral cell layer occurs in the external plexiform cell layer. In the mitral cell layer, the mitral cells are located. They form synapses in the inner plexiform layer with the granular cells of the granular cell layer. (B) HE staining of a sagittal section of the human olfactory bulb (b). Dashed box shows enlarged picture in (C). (C) Immunofluorescence staining of human olfactory bulb. Synaptophysin is shown in green. The specific layers of the olfactory bulb are marked. ONL: Outer nerve layer, GL: glomerular layer, EPL: External plexiform layer, MCL: Mitral cell layer, IPL: Internal plexiform layer, GCL: Glomerular cell layer. (D) Merged picture of ACE2 (red), OMP (white) and TUBB3 (green) illustrates the different layers of the olfactory bulb. High protein expression of TUBB3 and OMP can be found in the outer nerve layer, whereas ACE2 is mainly located in the glomerular layer, not co-localizing with TUBB3 or OMP. Nuclei are shown in blue, scale bar: 20 µm.

## DISCUSSION

In our present work we show that both virus entry proteins, ACE2 and TMPRSS2 are found in the sustentacular cells and Bowman’s glands of the olfactory epithelium (IV), but not in the primary sensory neurons. In the respiratory epithelium from the nasal septum (I), the intermediate nasal conchae (II), and the cellulae ethmoidales (III), ACE2 expression was detected in the basal cell layer and in the apical part of the respiratory epithelial cells as well as in the submucosal Gll. nasales. We conclude, based on our findings that SARS-CoV2 may bind to ACE2 and TMPRSS2 in epithelial cells of the respiratory, olfactory, and paranasal sinus epithelium and can thus penetrate the upper respiratory system. In the olfactory bulb, ACE2 was widely distributed with the highest expression in the glomerular layer, but was not co-localized with OMP positive neurons or other neuronal cell types.

Taken all findings together, this leads to the question, how the symptoms of anosmia can be explained in COVID-19 patients. Although it could be hypothesized that anosmia in COVID-19 is caused by viral infection of ORN, which further leads to their damage, and odorant information cannot be processed and projected to the brain via the olfactory bulb. However, in our work we can clearly demonstrate that there is no expression of ACE2 in the primary neurons, which supports results from previous work [9, 21]. Probably, the sustentacular cells are affected by the SARS-CoV2 virus, as these cells express both proteins ACE2 and TMPRSS2. Sustentacular cells are essential for the olfactory system. They do not only provide structural stability comparable to glia cells, but also support all other cells of the epithelium in a nutritious and metabolic way [14]. Moreover, sustentacular cells perform phagocytosis and are probably involved in protective mechanisms by expressing antiviral and antibacterial proteins [22]. Sustentacular cells are in close contact to the ORN, forming intercellular connections. Potentially, infected sustentacular cells may also invade ORN across these bridges, unassisted by the virus entry genes.

How does the localization of ACE2 and TMPRSS2 in Bowman’s gland cells and Gll. nasales of the respiratory epithelium fit to our hypothesis? Glandular cells in the nasal cavity, together with goblet cells, which are found in respiratory epithelia, are necessary for the production of mucus. Mucus is a viscous solution, which contains mainly water, ions, proteins and mucins and covers the apical side of the nasal epithelium [23]. It helps to maintain the physiological barrier of the epithelium against foreign substances from the air. In particular, the Bowman’s glands are supposed to produce enzymes providing xenobiotic-metabolizing functions [24]. Due to the virus entry protein expression in the Bowman’s gland and Gll. nasales, SARS-CoV2 could destroy many glandular cells which would probably result in a reduced or incorrect mucus production, leading to dysfunction or even damage of ORN. In addition, sustentacular cells and Bowman’s glands are supposed to produce odorant binding proteins (OBPs), which are indispensable for the perception of the odorants [25, 26]. Without OBPs in the mucus of the olfactory epithelium, the binding of odorants to the olfactory receptors is hampered.

That in mind, we hypothesize that a disruption of high numbers of sustentacular cells as well as mucus producing glandular cells could lead to a decreased perception of smell and leaves the olfactory epithelium less protected against other viral or bacterial threats.The stem cells of the olfactory epithelium, the basal cells, are most probably not affected by COVID-19 infections, maintaining the potential of reproducing ORN, sustentacular cells as well as Bowman’s glands [27]. This could explain the relatively fast recovery in most patients suffering COVID-19 triggered anosmia [28, 29]. However, some cases have been reported with very slow or nearly no recovery of anosmia [30]. Depending on the dimension of destruction of sustentacular and glandular cells, reproduction from basal cells followed by recovery of the nasal mucus may be prolonged or, as in severe cases, the destruction of sustentacular cells may also affect the basal cells leading to extended symptoms. In this context, it is noteworthy that we also found ACE2 expression in the basal cells of the respiratory epithelium.

Based on our findings we presume that SARS-CoV2 can enter the cells from the upper respiratory system via the viral entry proteins ACE2 and TMPRSS2. The infection of the sustentacular cells of the olfactory epithelium together with the underling Bowman’s gland cells may lead to altered mucus production, metabolism and structural instability in the olfactory epithelium. In addition, infection may result in the inability of the ORNs to connect to odorants via OBPs. Nevertheless, most patients regain their ability of smell perception, due to the fact that the basal cells are presumably not affected by the virus and can therefore replace destroyed cells of the olfactory epithelium.

## MATERIAL AND METHODS

### Ethics statement

Sampling of human material from body donors and all following experiments were made in accordance to local laws and regulations approved by the responsible ethical committee (Project number: 284/2020BO2).

### Tissue processing

Tissue was obtained from human body donors, with written consent. Surgery was performed transcranial with particularly attention to remove the cribriform plate together with the olfactory bulb as well as the nasal septum and the nasal conchae. Fixation of the whole specimen was performed for four days with daily change of Roti Histofix (Carl Roth, P087.1) fixation media, followed by one day washing step with PBS. Decalcification was achieved with 10 % EDTA for 70 days with medium change twice a week. After a washing step for one day with PBS, specimen was incubated for another day in 30 % sucrose, followed by embedding with Tissue-Tek (Thermo Fisher, 12351753) for frozen sections.

### Immunohistological staining

For Hematoxylin-Eosin staining 12 µm frozen sections were treated with filtered haematoxylin (Sigma, H9627) for 10 min, followed by a washing step with tap water for 2-5 min. Eosin (Sigma, 230251) counterstaining was performed for 7 min, followed by careful rinses with distilled H_2_O and dehydration with increasing alcohol concentrations (70 %, 95 %, 100 %). After washing two times in Xylene, sections weremounted with Depex mounting media (VWR, 13514).

### Immunofluorescence staining

For Immunofluorescence staining 12 µm frozen sections were rehydrated for 5 min with PBS (Thermo Fisher, 10010056) followed by an ethanol gradient each 30 sec (70 %, 95 %, 99 %, 95 %, 70 %). Frozen sections were blocked in skimmed milk blocking buffer (TBS + 10 % NDS + 1,25 % BSA, 4 % skimmed milk, 0,1 % Triton-X) for 30 min at room temperature. Primary antibody were diluted in skimmed milk blocking solution and incubated over night at 4 °C. The following primary antibodies were used: ACE2 rb (Abcam, ab15348 1:100), TMPRSS2 ms (Santa Cruz, Sc515727, 1:50), TUBB3 ms (BioLegend, 802002, 1:1000), OMP gt (WAKO, 544-1001 1:500). Synaptophysin rb (Abcam, 14692-100, 1:100). All primary antibodies were verified in appropriate tissue (data not shown). After several washing steps, secondary antibodies were diluted 1:100 in PBS together with DAPI (Abcam, ab228549) and incubated for 45 min in room temperature free of daylight. Following secondary antibodies were used: Dαrb 488 (Invitrogen, A32790), Dαms 488 (Invitrogen, A32766), Dαrb 546 (Invitrogen, A10040), Dαgt 546 (Invitrogen, A11056), Dαms 647 (Abcam, ab150107). Section were embedded with Mowiol (Carl Roth, 0713). Immunofluorescence stainings were analysed using the Axio Imager. M2 microscope with the AxioVision software (Zeiss).

Scans of immunolabelled whole nose specimen were done by order of Zeiss, pictures were analysed with ZEN Blue software (Zeiss).

## AUTHOR CONTRIBUTIONS

M.K., S.K., A.M. invented and designed the project. A.M., P.H.N. and A.W. performed the surgeries and extraction of human tissues. A.M. and M.K. performed the tissue processing procedures and generated the frozen sections. A.M. performed the histological stainings. S.K. optimized and performed the immunofluorescence stainings. M.K. took and edited the microscope pictures and created the Figures and schemata. S.K. and M.K. wrote the manuscript. A.W., A.F.M., P.H.N. helped discussion the project and performed proofreading of the manuscript. M.K., A.K. and S.L. were proofreading of the manuscript. Financial support was provided by S.L.

